# Large-scale discovery of potent, compact and lineage specific enhancers for gene therapy vectors

**DOI:** 10.1101/2023.10.04.559165

**Authors:** Nikoletta Psatha, Pavel Sova, Grigorios Georgolopoulos, Kiriaki Paschoudi, Mineo Iwata, Jordan Bloom, Tatyana Ulyanova, Hao Wang, Alexandra Kirtsou, Ninos-Ioannis Vasiloudis, Matthew S. Wilken, John A. Stamatoyannopoulos, Evangelia Yannaki, Thalia Papayanopoulou, George Stamatoyannopoulos, Jeff Vierstra

## Abstract

Regulation of gene expression during cell development and differentiation is chiefly orchestrated by distal noncoding regulatory elements that precisely modulate cell selective gene activity. Gene therapy vectors rely on the cellular and context specificity of regulatory DNA elements to express therapeutic transgenes in the correct location and time. Here, we develop a straight-forward, one-shot approach to screen putative regulatory sequences identified in large-scale epigenomics profiling experiments for precise and programmable control of transgenes encoded within gene therapy viral vectors. We designed a library of 15,000 short sequences (∼200bp) derived from a set of developmentally active DHS elements during human *ex vivo* erythropoiesis and cloned them into a GFP reporter lentiviral vector. In an erythroid progenitor cell line, these elements display a gradient of transcriptional enhancer activity, with some demonstrating equivalent activity to the canonical β-globin μLCR despite a 9-fold smaller size. We show that these elements are both highly cell type restricted and developmental stage specific both *in vitro* and *in vivo*. Finally, we replace the μLCR element with one of the novel short enhancers in a β-thalassemia lentiviral therapeutic vector and efficiently correct the thalassemic phenotype in patient-derived HSPCs. More broadly, our approach provides further insights into enhancer biology with wider implications into the development of highly cell type specific and efficacious viral vectors for human gene therapy.

## Introduction

Gene therapy has heralded a fundamental shift in modern medicine and has enabled curative single dose treatments for an array of diseases. The overall goal of this therapeutic modality is to achieve durable expression of a therapeutic gene or transgene that ameliorates or eliminates molecular disease symptoms. Conventional gene therapy viral vectors consist of two main components: 1) therapeutic transgene and 2) a regulatory sequence driving the transgene expression. For several genetic or acquired diseases, gene therapy has successfully moved from proof-of–concept studies to clinical translation and commercially marketed clinical therapies^1–3^. Despite this promise, development of new gene therapy-based treatments has proven challenging due to transgene expression failing to reach therapeutic levels and substantial off-target effects (i.e. expression of the transgene in non-target cell types and/or ectopic activation of oncogenes near viral integration sites). Many of these limitations can be directly linked to the architecture of gene therapy vectors. A straight-forward strategy that can simultaneously amplify efficacy and safety of transgene vectors is the utilization of cell-type specific, compact regulatory sequences in the vector designs to precisely control expression of therapeutic transgenes in their proper context.

Recently, genome scale chromatin accessibility assays have identified millions of putative *cis*-regulatory elements^4,5^. The vast majority of these potential *cis*-regulatory elements are cell-selective and developmentally specific and, thus in principle, can be harnessed to control therapeutic transgene expression in a wide variety of contexts (i.e., location and time). Despite systematic efforts to biochemically delineate human regulatory DNA^6–8^, characterizing regulatory function (i.e., impact on gene expression) has remained challenging. While massively parallel reporter assays (MPRAs) can identify sequences with regulatory function, in many cases these assays have limited translatability because they are performed using non-integrating episomal vectors in highly proliferating malignant cell lines and not the therapeutically relevant cell type^9–11^. Moreover, when integrating vectors are employed (lentiviral), the readout of the assay is not a transgene, but the transcription of the regulatory sequence itself (located 3’ to the promoter). In this case, the architecture of the screening vector does not resemble a putative therapeutic (i.e., transgene control). Taken together, these limitations urge the development of alternative enhancer screening methods to identify and design control sequences for gene therapy vectors.

In this study, we describe a direct enhancer discovery pipeline to optimize gene therapy vectors including identification of cell type specific enhancer elements and design of a new therapeutic vector. As a proof of concept, we apply this approach to β-hemoglobinopathies, the most common monogenic disorders worldwide and demonstrate that novel, cell type specific, compact, potent enhancers challenge the prototypic μLCR enhancer (a truncated version of the β-globin locus control region) of hemoglobinopathy vectors and enable efficient phenotypic correction.

### Large-scale evaluation of erythroid enhancers in a chromosomal and therapeutically relevant context

We previously mapped all developmentally regulated DHSs (*n*=11,805) during human erythropoiesis^12^. Here, we focused on the *de novo* activated elements during erythroid differentiation. We excluded DHSs accessible at CD34^+^ hematopoietic stem and progenitor (HSPC) stages to avoid elements potentially active in other lineages. Fully, this resulted in 5,393 DHSs accessible at different stages of erythroid differentiation where 80% of the DHSs exhibit peak accessibility after day 7 of *ex vivo* erythroid differentiation (**Supplementary** Figure 1a). In their majority, these DHSs represent distal non-coding elements either intronic or intergenic or (49.7% and 30.9%, respectively) whereas promoters are relatively under- represented (11.4%) (**Supplementary** Figure 1b). As expected, these elements display strong lineage-specific accessibility as DNase I signal in non-erythroid hematopoietic cell types, including CD34^+^ hematopoietic stem and progenitor cells (HSPCs) is markedly low (**Supplementary** Figure 1c).

To functionally screen our candidate regulatory elements (CREs), we designed a lentiviral vector closely matching those currently used in clinical therapeutics^1,2,13^. Our screening method deviated from traditional lentivrial MPRAs by employing a modified FACS based lenti-MPRA approach where the evaluation of the CREs was based on their effect on protein expression (**Figure 1a**). The enhancer screening library generated from tiling each of the 5,393 erythroid DHSs (median size 233bp) into overlapping 198bp-long oligos, resulting in a library of 14,999 DHS fragments with a median of 3 tiles per DHS (**Figure 1a** and **Supplementary** Figs. 2a,b). The DHS-oligo library was cloned upstream of a minimal (169bp) human β- globin promoter driving GFP expression (**Figure 1b**). To avoid positional effects driven by random vector integration we included recently identified, dual functioning (enhancer-blocker and barrier) human chromatin insulator into our lentiviral vector which flanked our expression cassette^14^. To maintain the cell context as close to clinical translation as possible, we performed screening a CD34^+^ cell derived human erythroid progenitor cell line, HUDEP-2 cells, that roughly corresponds to day 7 adult erythroid progenitors on the basis of chromatin accessibility^12^ (**Supplementary** Figure 3). Critically, we performed all HUDEP- 2 transductions at multiplicity of infection (MOI) of 0.4 to achieve a theoretical maximum of one viral integration per cell.

**Figure 1.**
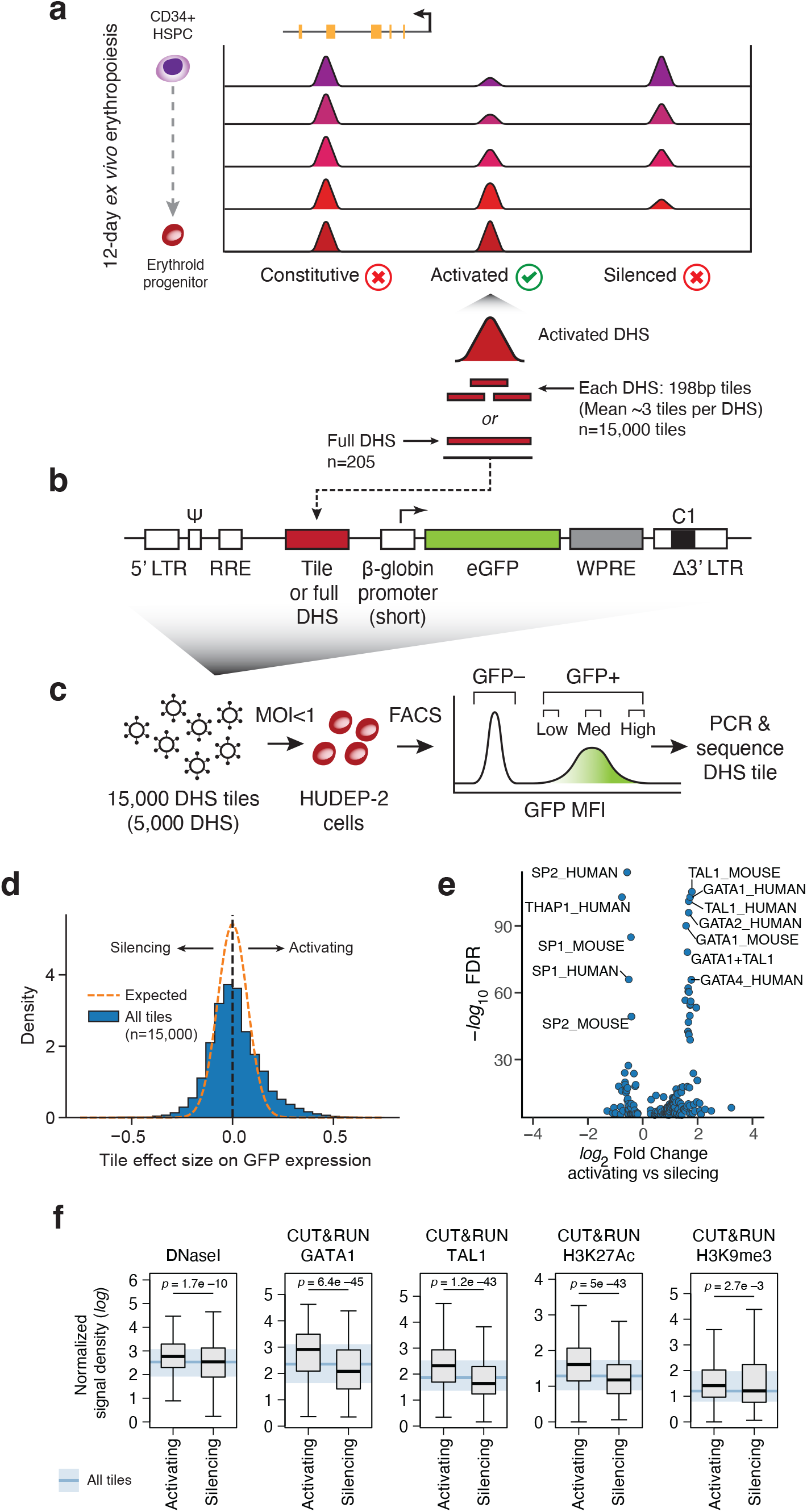
Selection and screening of putative erythroid enhancers. **(a)** Strategy for selecting candidate erythroid DNA regulatory sequences for enhancer screening. 5,393 DNase I hypersensitive sites (DHS) activated during *ex vivo* erythropoiesis were selected and ∼3 198bp- long tiles per DHS were synthesized to construct a screening library of 15,000 sequences. In the second screening round, 205 full-length DHS were synthesized. **(b)** The library was cloned into a lentiviral vector upstream of a minimal β-globin promoter, driving the expression of eGFP, and the C1 chromatin insulator in the 3’ LTR. **(c)** HUDEP-2 cells were transduced at a MOI<1 and sorted by flow cytometry into discrete bins of varying GFP intensity (low, medium, high). **(d)** Histogram depicting the distribution of the maximum likelihood estimated effect of each tile on GPF expression. Dashed line represents the expected null distribution. **(e)** Volcano plot of the differentially enriched transcription factor binding motif archetypes between the top 1000 silencing (log_2_ fold change < 0) and the top 1000 activating tiles (log_2_ fold change > 0). **(f)** Differences between the top 1000 activating and top 1000 silencing tiles in terms of chromatin accessibility by DNase I, GATA1 and TAL1 occupancy as well as enrichment for two chromatin marks as measured in HUDEP-2 cells. Boxplot whiskers extend to 1.5X IQR. Kruskal-Wallis *p*-values are reported.

In vectors containing only a promoter, GFP expression was low and the overall population was partially overlapping with the untransduced/non-GFP expressing cells (**Supplementary** Figure 4a). Five days post transduction we FACS-sorted the transduced cells based on GFP intensity (low, medium and high), sequenced the integrated lentiviral cassette, and computed the relative enhancer tile frequencies in each GFP bin (**Figure 1c and Supplementary** Figure 4b). Because expression of integrated lentiviral cassettes is impacted by local genomic and epigenomic context (i.e., position-effect variegation)^15^, we assessed the number of unique integrations required for each element to yield reliable expression estimates. We found that for a library of our size (∼15,000 enhancer tiles), ∼800 integrations were need per-element to achieve robust replicate concordance (*r*>0.9), and that lower number of integrations resulted in poor replicate concordance (100X *r*=0.347; and 200X *r*=0.423 respectively) (**Supplementary** Figure 5), highlighting that lentiviral MPRA approaches are critically dependent on the number of unique insertions to overcome positional variegation effects^15^.

At a library coverage of 800X, we recovered virtually all of the original enhancer tiles (14,668 fragments in total, 97.8% of the designed library). To estimate the effect of each fragment on altering GFP intensity we employed a statistical framework (adapted from ^16^), that estimates the latent effect each sequence on expression by modeling that the tile frequencies in each GFP bin through maximum likelihood. Using this ML approach, we ranked each enhancer tile based on its estimated effect value, resulting in 6,577 activating elements and 8,091 silencing elements (**Figure 1d**). The latter suggests that a large number of DHSs activating during erythroid differentiation may function as repressors.

Enhancer tiles strongly associated with high GFP expression were marked by increased DNase I accessibility, binding of master erythroid regulatory factors (e.g., GATA1 and TAL1) and histone post-translational modifications canonically associated with enhancing *cis*-regulatory elements (H3K27Ac, H3K9me3) (**Figure 1d**). To assess how the full-length DHSs corresponds to the respective tile in terms of enhancer activity, we randomly selected a subset of 202 elements (**Supplementary Table 1)** from the overall activating elements (*n*=6,577). For each element in this smaller library we synthesized a new sequence element corresponding to the encompassing full-length DHS for each tile and observed the effect of the strongest tile from each DHS on GFP expression to be well-correlated (*r*=0.51) to that of the full-length DHS (**Supplementary** Figure 6), indicating that for many DHS elements, a compact sub-sequence encodes the majority of its regulatory function.

### Identification of enhancers with lineage-selective and tuned developmental activity

Proper cell lineage and temporal activity of transgenes is paramount to safe and effective gene therapy vector. To test for lineage specificity and developmental temporal activity of these sequences we individually characterized a subset of 40 candidate DHSs from the mini-library (**Supplementary Table 2**). We cloned each one individually (in triplicate) in the same screening vector and finally compared against the HS2 and HS3 of the β-globin LCR, and a no-enhancer vector and then transduced them into HUDEP-2 cells to evaluate the enhancer activity of the individual elements. We found that 38 (95%) had a marked increase in GFP expression vs. the no enhancer control vector (**Figure 2a**). Critically, the expression intensity values from the individual vectors were well correlated with the activities measured from the mini-library (*r*=0.72, **Supplementary** Figure 7).

**Figure 2.**
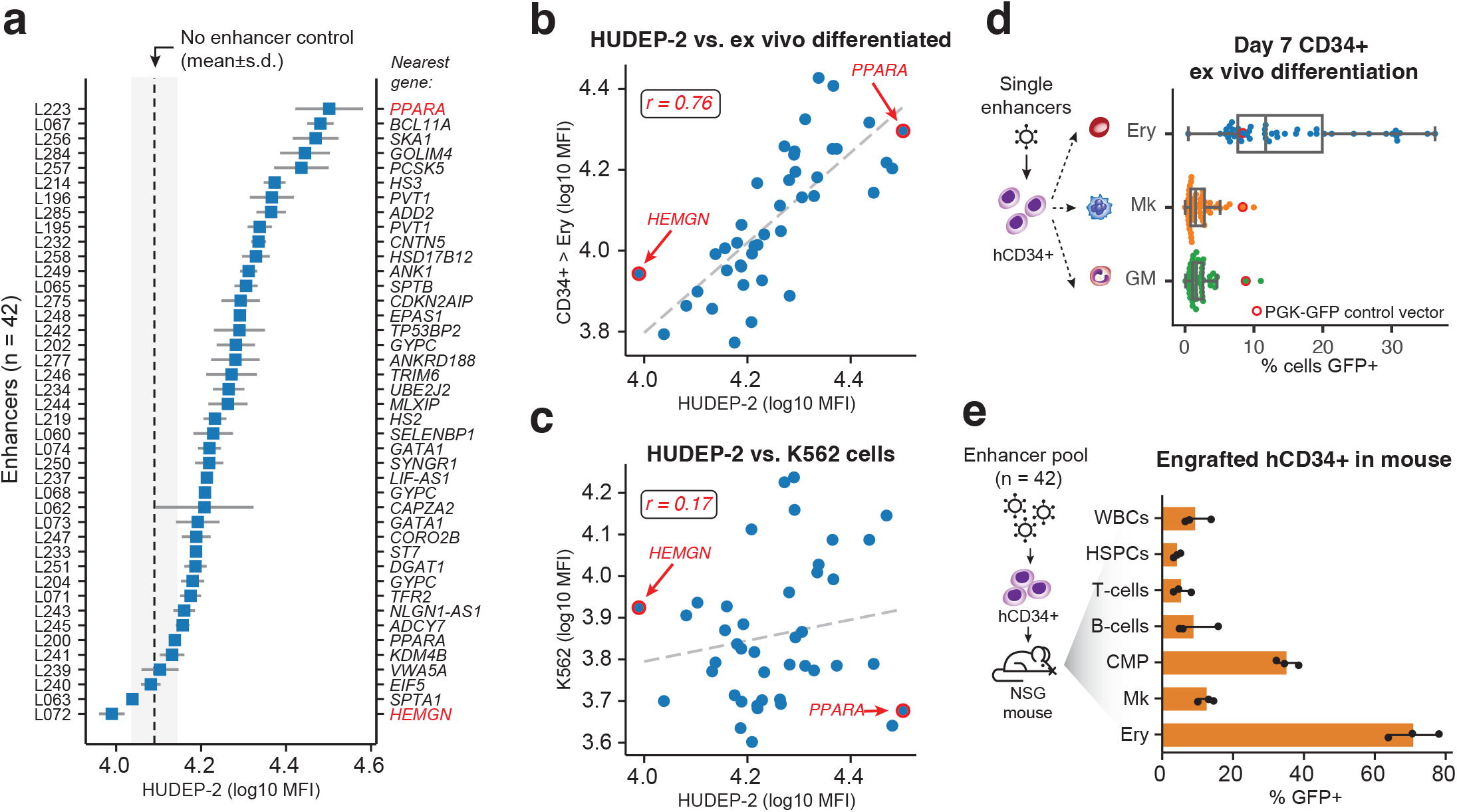
Novel enhancer elements display erythroid-specific activity in vivo and in vitro. **(a)** GFP expression of the 40 DHSs individually cloned in the same backbone vector and transduced in HUDEP2 cells at a MOI<1. Full length β-globin HS2 and HS3 were used as positive controls and a no- enhancer vector as negative control. Mean±SD is shown. **(b)** Correlation between the GFP intensity (log_10_ MFI) of the 42 selected elements after transduction in HUDEP-2 cells (*x*-axis) and in CD34^+^ differentiated erythroid cells (y-axis). Pearson’s r correlation coefficient is shown **(c)** Correlation between the GFP intensity (log_10_ MFI) of the 42 selected elements after transduction in HUDEP-2 cells (x-axis) and in K562 cell line (*y*-axis). Pearson’s *r* correlation coefficient is shown. **(d)** Each of the 42 identified enhancers is individually transduced into CD34^+^ cells and each pool (*n*=42) is subjected to *ex vivo* erythroid (Ery), megakaryocytic (Mk) and granulocytic-monocytic (GM) differentiation. After 7 days of differentiation the percentage of GFP^+^ cells are determined by flow cytometry. Boxplot whiskers extend to 1.5X IQR. The cell cultures transduced with the PGK-GFP control vector are highlighted in red. **(e)** Mobilized peripheral blood CD34^+^ cells from healthy donors were transduced with a lentiviral library of the identified 42 vectors. The cells were transplanted into NBSGW mice and bone marrow was collected 16 weeks post transplantation. Percent of GFP^+^ cells was assessed by flow cytometry in all engrafted human hematopoietic lineages.

We then performed parallel experiments in adult human CD34^+^-cell derived erythroid cells which represent the target cell type for *ex vivo* human gene therapy and in the erythroleukemia K562 cell line, which has long served as a popular erythroid cell model. Notably, the activity of the enhancers in K562 cells was poorly correlated to HUDEP-2 cells (*r*=0.23) (**Figure 2b**). By contrast, the same enhancer constructs transduced into primary mobilized CD34^+^ HSPCs followed by *in vitro* erythroid differentiation (7 days, erythroblast stage) displayed activities that closely followed that of HUDEP-2 (*r*=0.82) (**Figure 2c**). This result emphasizes the importance of cell type context in screen assays for the functional interrogation of regulatory DNA sequences.

As mentioned above, all tested DHSs were selected based on their increased accessibility in adult erythroid cells. However, this feature alone does not exclude a potential activity of these elements in other hematopoietic cell lineages. This is particularly relevant to the development of gene therapy products where transgenes integrating near proto-oncogenes have the potential to drive malignant transformations due to lineage-promiscuous enhancer elements. To assess to what degree the activity of these enhancers are restricted to the erythroid lineage, we transduced mobilized CD34^+^ HSPCs with each vector individually and allowed them to differentiate (*ex vivo*) towards erythroid, myeloid (granulocytic/monocytic) and megakaryoctyic lineages (**Figure 2d** and **Methods**). For nearly all enhancers we found robust GFP expression within the erythroid lineage (mean 15% GFP^+^), while minimal-to-no expression was observed in the other lineages (mean 2.3% GFP^+^ for each megakaryocytic and myeloid). A generic PGK-GFP vector was used as a positive control which was expected to drive GFP expression uniformly across all lineages (**Figure 2d**). Only one of the tested enhancers (an element proximal to *SKA1*) displayed activity in all cell lineages and upon investigation the site was found accessible across the myeloid lineages. This result provides strong evidence that in the current system lineage specificity is driven exclusively by the enhancer and independent of the promoter.

To validate these findings *in vivo* and expand beyond the myeloid lineages, we performed xenotransplantation experiments where human CD34^+^ cells were transduced with a pool of the described vectors. The transduced cells were transplanted in the NBSGW mouse model which efficiently supports multilineage human hematopoiesis. Measurements at 16-weeks post-transplantation revealed robust GFP expression restricted to human erythroid progenitors (70% GFP^+^ in hCD45^+^/CD235a^+^) and myeloerythroid progenitors (38% GFP^+^ in hCD45^+^/CD33^+^), with negligible levels of GFP detected within the human, non-erythroid compartments (**Figure 2e**). Cumulatively, individual evaluation of the candidate sequences revealed graded activity, enabling the precise modulation of transgene expression, in both amplitude and lineage specificity.

### Enhancer-driven transgene activity parallels temporal dynamics of chromatin accessibility

The DHS elements included in the enhancer library were selected for their dynamic accessibility profile in the course of *ex vivo* erythroid differentiation^12^. We investigated whether these elements regulatory capacity (i.e., effect on transcription) paralleled the accessibility profile within their native genomic context. We transduced individual pools of CD34^+^ cells, each with one of the 40 selected enhancer elements and subjected them to *ex vivo* erythroid differentiation, and for each culture we measured GFP MFI as a proxy for enhancer function at regular intervals during the differentiation (**Figure 3a**). Comparing the GFP MFI with the accessibility of their corresponding DHS during differentiation we found a strong correlation between the two across the elements tested (median *r*=0.79) (**Figure 3b**) strongly indicating that temporal enhancer functionality is maintained outside of their native genomic context and is solely encoded by the local sequence (∼200bp) of these elements.

**Figure 3.**
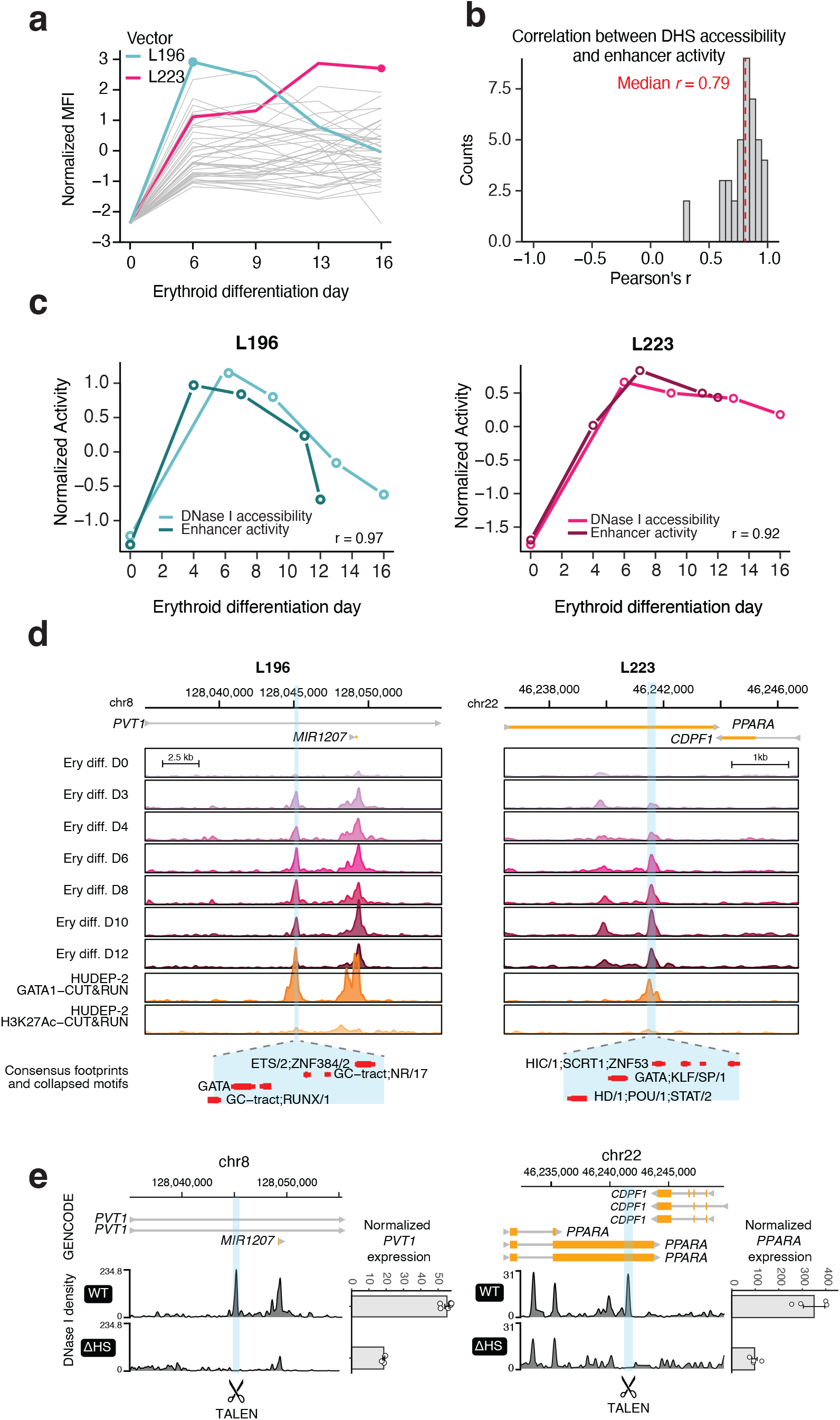
Novel enhancer elements are native transcriptional enhancers with erythroid temporal activity. **(a)** We assessed the enhancer activity kinetics of the 42 elements by measuring GFP expression with flow cytometry (MFI, *y*-axis) during *ex vivo* erythroid differentiation from CD34^+^ cells. An early active (L196, teal) and a late active (L223, magenta) element were selected for further study. **(b)** Correlation (Pearson’s *r*) between the temporal profiles of enhancer activity (MFI) of the 42 elements and their *in situ* DNase I accessibility profile during *ex vivo* erythropoiesis. Median *r* is denoted with a dashed red line. **(c)** Comparison between the enhancer activity (MFI) and in situ DNase I accessibility for the L196 vector (left) and L223 (right). **(d)** The enhancer element in the L196 vector is derived from a DHS intronic to the *PVT1* lncRNA and the element in L223 is an exonic DHS in the *PPARA* locus. Signal density tracks of the DNase I accessibility during erythroid differentiation, GATA1 CUT&RUN occupancy and H3K27Ac CUT&RUN enrichment in HUDEP-2 cells is shown for *the PVT1* locus (left) and *PPARA* locus (right). Bottom track shows consensus footprinted motifs overlapping with each element. **(e)** Genetic deletion experiments of *PVT1* DHS and *PPARA* DHS in HUDEP-2 cells result in significant repression of *PVT1* and *PPARA* expression, respectively. Browser tracks of the DNase I accessibility in WT HUDEP-2 and mutant (ΔHS) are shown. Barplots show the mean±SE of normalized gene counts from *n*=4 experiments.

Given that the enhancers display a precise temporal activity we then interrogated the *in situ* functionality of strongest enhancers: L196 (early activation) and L223 (terminal activation) (**Figure 3c**). Surprisingly, despite their strong enhancer function in both HUDEP-2 and *ex vivo* differentiated adult HSPCs, the two elements display weak H3K27ac levels (**Figure 3d**), a histone modification typically associated with active enhancers. Despite the limited presence of H3K27ac, homozygous knockout these elements (**Supplementary** Figure 8a) resulted in strong, reproducible transcriptional repression of their target genes (*PVT1* for L196 and *PPARA* for L223) (**Figure 3e and Supplementary** Figure 8b). These results strongly indicate that these enhancers mirror their developmental activity in an engineered ectopically integrated vector and can therefore be used for temporal control of transgene expression in a gene therapy setting.

### Direct preclinical translation of the novel enhancers

Because these enhancers drive robust gene expression, and their activity reflects observed developmental dynamics, we sought to apply them directly to a therapeutic vector for β-hemoglobinopathies as a translational paradigm. Here, we focused on the *BCL11A*-shRNA transgene approach. First, *BCL11A* knock down leads to HbF reactivation which is easily measurable in both healthy and diseased human erythroid cells and in addition. Secondly, *BCL11A* knock out in any other than the erythroid lineage would have an immediate negative effect in cells’ viability, proliferation, engraftment and differentiation capacity *in vivo*. Therefore, the new shRNA vectors incorporating the optimal erythroid enhancers would permit us to simultaneously test their lineage-specificity and by extension, safety as well as their efficiency. A *BCL11A*- shRNA^13,17,18^ was cloned in the same backbone as the screening vector (**Figure 4a**). Of note, both the *PPARA*–shRNA and *PVT1*–shRNA vectors showed a significant increase in viral production compared to the µLCR vector, likely due to the small size of the new enhancers (∼9 times smaller than the clinically utilized µLCR) (**Figures 4a,b**). Subsequently, all shRNA vectors were transduced in HUDEP-2 cells, to evaluate HbF reactivation in a cell line with a low background of γ-globin expression. The small size of the new enhancers did not reduce their therapeutic effect as they were both able to achieve γ-globin reactivation in transduced cells at the same levels as the μLCR vector (**Figure 4c**). HbF expression in untransduced cells (GFP-) was maintained at low levels, as expected. Subsequently, we transduced CD34^+^ cells from healthy donors with the above vectors with serial dilutions of the viruses which showed that the significantly larger μLCR-shRNA vector quickly reaches a plateau in HSPCs transduction ability (**Supplementary** Figure 9a,b). HbF expression within the transduced populations was similar or higher using the new vectors compared to the μLCR-shRNA, with *PVT1-*shRNA achieving overall the highest percentage among the three groups (**Supplementary** Figure 9c), while the differences in HbF expression were not dependent to the MOI used (**Supplementary** Figure 9d).

**Figure 4.**
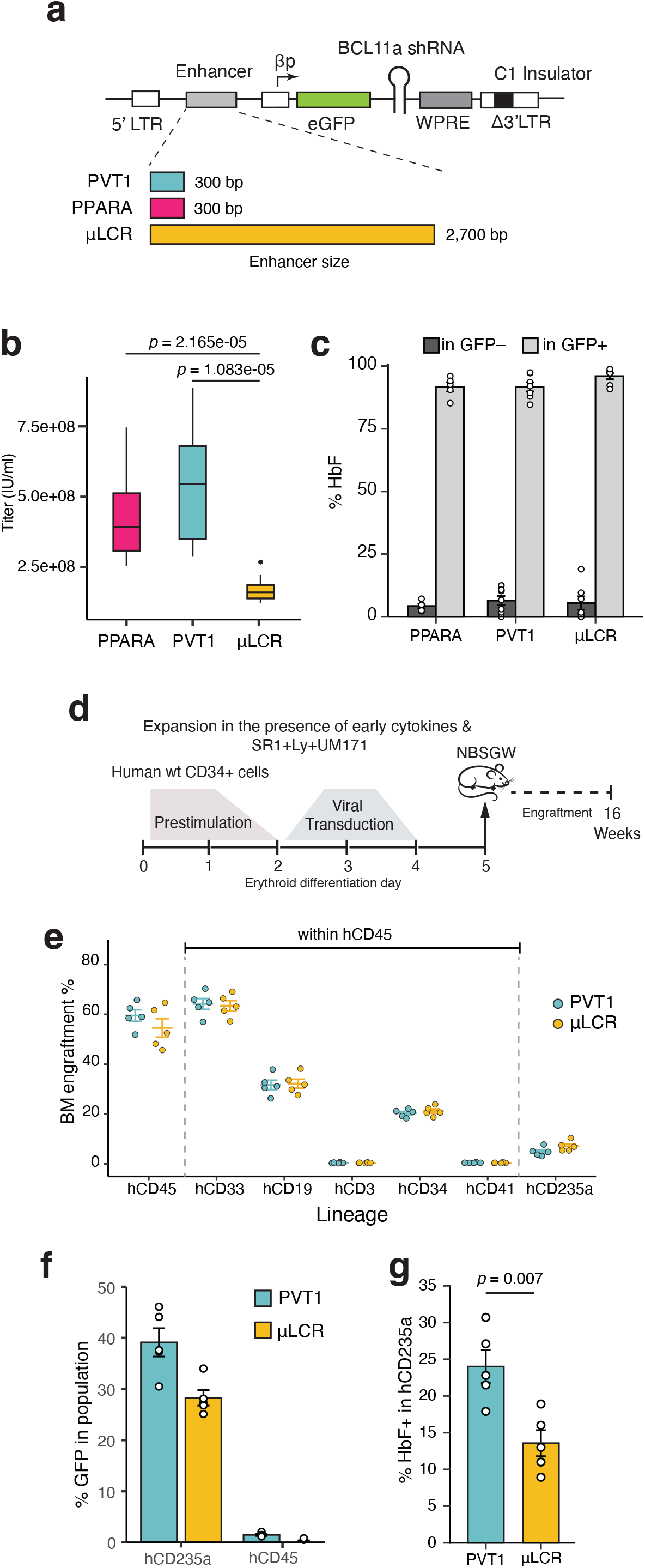
shRNA vectors equipped with the novel enhancers outperform μLCR in vitro and in vivo. **(a)** Schematic representation of the vector design showing the difference in size of the new enhancers compared to the μLCR. C1: chromatin insulator, βp: minimal β-globin promoter. **(b)** Titration of the *PPARA*, *PVT1* and μLCR BCL11A-shRNA vectors in K562 cells based on qPCR. **(c)** HbF expression (%) within the GFP- and GFP^+^ populations assayed by flow cytometry in transduced HUDEP-2 cells with one of the *PPARA*, *PVT1*, and μLCR *BCL11A*-shRNA vectors. **(d)** Workflow of the *in vivo* experiments. Mobilized peripheral blood CD34^+^ cells from healthy donors were transduced at the same MOI with the μLCR and *PVT1*-shRNA vectors in the presence of cytokines and a small molecule cocktail (SR1: StemRegenin 1, Ly:Ly2228820, UM171). Two days post transduction an equal number of cells was transplanted in NBSGW mice. Mice were sacrificed 16 weeks post transplantation. **(e)** Human chimerism in the bone marrow of the transplanted mice. hCD33, hCD19, hCD3, hCD34, hCD41 subpopulations were calculated within the hCD45 population. **(f)** GFP expression (as %GFP^+^ cells) by flow cytometry in erythroid (hCD235a^+^) and non-erythroid (hCD45^+^) engrafted cells. **(g)** *In vivo* HbF expression (as % HbF+ cells) in engrafted human erythroid (hCD235a^+^) cells at the time of sacrifice.

To assess the new vectors performance *in vivo*, we transduced at the same MOI normal mobilized peripheral blood derived CD34^+^ cells with the *PVT1*-shRNA vector and compared its effect with that of the μLCR- shRNA vector in a xenotransplantation mouse model (**Figure 4d**). Transduction of the infused cells was performed at the same MOI between the two vectors. The mice were sacrificed 16 weeks post transduction and their bone marrow was analyzed for multilineage reconstitution and HbF expression (within the erythroid lineage). Importantly, multilineage reconstitution was observed with the *PVT1* enhancer (**Figure 4e**). As expected from our previous experiments, GFP expression was restricted to the erythroid compartment, with higher number of transduced cells in the *PVT1* group despite the cells being transduced with the same MOI (**Figure 4f**). The highest transduction/transgene expression rate was also translated to a higher overall percentage of HbF^+^ cells in the *PVT1* group compared to the LCR (**Figure 4g**).

### *Ex vivo* therapeutic correction of β-thalassemia major in primary, patient-derived HSPCs

Given the results from transduced healthy HSPCs we next tested our enhancers in cells derived from patients with β-thalassemia major (**Supplementary Table 2**). Expanded HSPCs derived from two mobilization clinical trials^19,20^ were transduced with the *PVT1* and μLCR-shRNA vectors (**Methods** and **Supplementary** Figure 8) and cultured in erythroid medium for 18 days (terminal differentiation). Recapitulating the results previously observed in normal HSPCs, GFP transduction rates for the *PVT1*- shRNA vector was significantly higher compared to the μLCR (*p*=0.04, **Supplementary** Figure 9). Furthermore, we observed that the *PVT1*-shRNA vector had markedly increased proliferative ability at later stages of differentiation (>3-fold increase in total erythroid cells on day 1; *p*=0.0015 vs. untransduced) (**Figure 5a**). A similar trend but to a lesser degree (2.6-fold), but without statistical significance, was observed for the μLCR-shRNA transduced cells (*p*=0.1 vs untransduced). HbF^+^ cell frequency in the untransduced population was surprisingly high, which is compatible with a survival advantage of cells with elevated HbF levels. However, despite the high background, HbF expression in both transduced groups was significantly higher in the enucleated and nucleated populations compared to the untransduced cells, with the difference being higher in the case of *PVT1* compared to that of the μLCR (*p*=0.003 vs. *p*=0.019 vs untransduced, **Figure 5b**). Similarly, transduction increased production of both Aγ and Gγ chains in both groups (α/γ-globin: *p*=0.024 and *p*=0.36, μLCR and *PVT1* vs. untransduced) and restored the α:β-like chain balance to 1:1 (**Figure 5c,d**).

**Figure 5.**
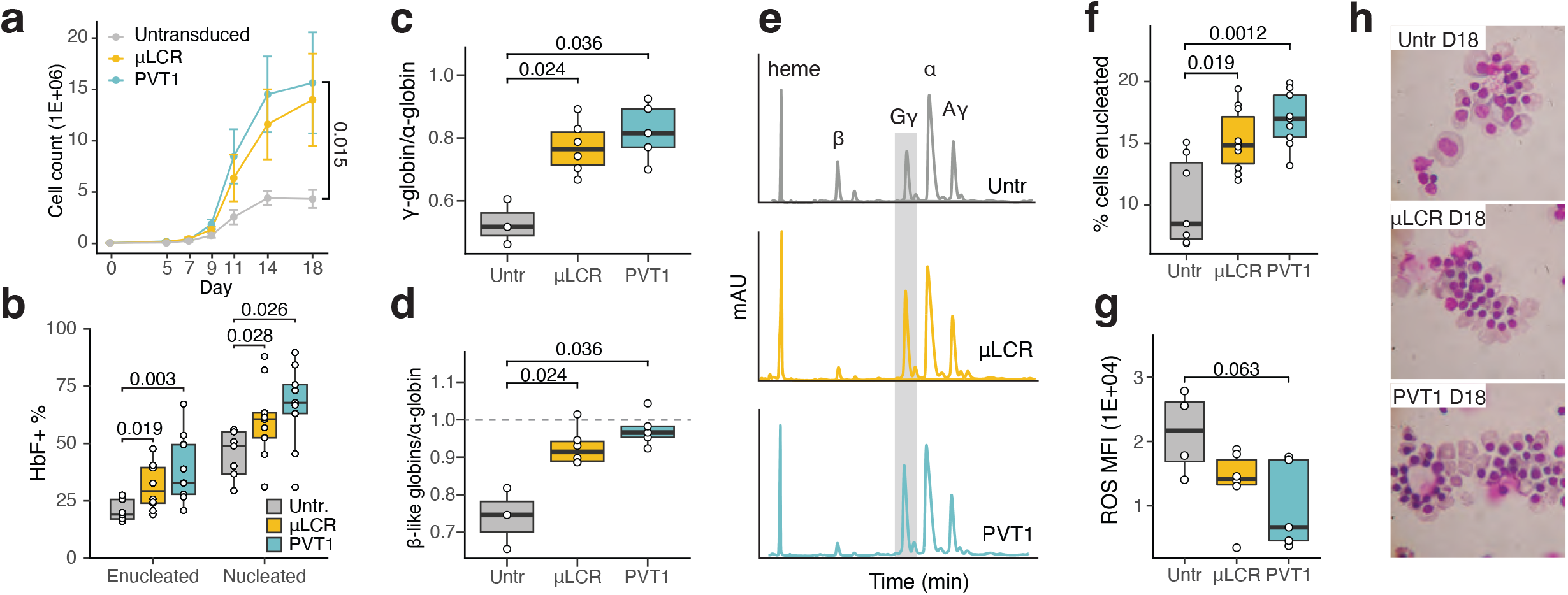
Therapeutic vectors equipped with PVT1 enhancer achieve superior correction of the β- thalassemia phenotype compared to μLCR. Mobilized peripheral blood CD34^+^ cells from 5 patients with beta-thalassemia major were transduced with the PVT1 and μLCR-shRNA vectors in the presence of cytokines and small molecules. The cells were cultured under erythroid conditions for 18 days, 2 days post transduction. **(a)** Absolute cell numbers during erythroid differentiation of transduced and untransduced cells. **(b)** HbF expression (as %HbF^+^ cells) measured by flow cytometry in enucleated and nucleated cells on day 18 of *ex vivo* erythroid differentiation. **(c)** Globin chain production displayed as γ:α and total β-like:α ratio in transduced and untransduced cells measured by HPLC. **(d)** HPLC trace profiles of globin chains in transduced and untransduced β0/β^+^ patient CD34^+^ derived erythroid cells. **(e)** Percentage of enucleated cells on day 18 of the differentiation measured by flow cytometry. **(f)** Flow cytometry measured intensity of reactive oxygen species (ROS) post CFSE staining of erythroid cells on day 11 of the erythroid culture. **(g)** Characteristic morphology of transduced and untransduced erythroid cells in different erythroid differentiation and maturation stages.

This increase resulted in improved enucleation of the transduced- over the untransduced cells (Figure 5e), more pronounced in the *PVT1*-shRNA-group compared to μLCR-shRNA-group, (*p*=0.0012 and *p*=0.019, respectively) that was accompanied by a marked improvement of the thalassemic phenotype in transduced populations. Specifically, higher numbers of maturing erythroid cells at the end of EC were observed in the transduced cells, in contrast to an erythroid maturation blockade in the untransduced population as well as reduction of the high levels of reactive oxygen species (ROS), a hallmark of oxidative stress in thalassemia caused by either increased autoxidation of α-globin chains or by increased iron levels (**Figure 5f, g**).

Overall, our results indicate that this new generation of vectors incorporating novel enhancers have advantages over conventional vector designs thus overcoming long-standing limitations of viral vectors, and that these compact, powerful and erythroid-specific enhancers can successfully replace the β-globin μLCR, the gold standard so far in globin vector development.

## Discussion

Gene therapy has recently met great success for a variety of monogenic hereditary diseases, cancer, and neurodegenerative disorders and holds great promise in becoming a standard durable clinical option for many disorders of genetic or epigenetic background. Nonetheless, current vector designs suffer by suboptimal safety and efficacy in terms of tissue specificity, transgene expression levels, and viral titers^21^. *In vivo* gene therapy in particular, requires both tissue specificity and tightly controlled gene expression as the use of strong promoter-enhancer pairs can result in tissue toxicity due to transgene overexpression^22–24^. In the context of ex vivo gene therapy where high or even supranormal therapeutic protein levels are required, transgene expression is driven by strong promoter^2,25^ or enhancer sequences^26,27^. Unfortunately, a lentiviral vector equipped with the constitutive MNDU3 promoter developed for cerebral adrenoleukodystrophy (CALD) gene therapy lead to the first recorded genotoxic events as three pediatric patients developed myelodysplastic syndrome (MDS) and/or clonal dominance^28^. In addition to adequate transgene expression levels, successful gene therapy protocols mandate high transducibility and efficient viral production. Hemoglobinopathies, a popular disease model for gene therapy has a history of vector development for more than 30 years and a recently approved product for β-thalassemia. The incorporation of the μLCR enhancer has enabled both therapeutic transgene expression levels and erythroid-specific expression patterns, representing the *sine qua non* in globin vector design. However, its large size (∼3kb) yields low viral titers rendering vector production suboptimal and highly expensive in addition to impeding efficient cell transducibility^29,30^.

Here, we introduce a platform for large-scale discovery of short and potent enhancer elements with precise spatiotemporal activity, enabling the development of a new generation of viral vectors towards efficient, safe, and cost-effective gene therapy applications. Capitalizing on our previous efforts of systematic mapping and characterization of the cis regulatory DNA elements involved in human erythropoiesis^12^, we designed a flow cytometry-based screening strategy using a lentiviral vector that allowed us to assess the transcriptional enhancer activity of 15,000 short sequences.

Although large scale lentiviral screening strategies have also been developed by others^31^, these approaches largely rely on highly-proliferative cell lines^31–36^ and/or non-human models^37–40^. It is now well understood that the activity of c*is*-regulatory elements is determined by their sequence composition, genomic context, and the availability of trans-acting proteins (transcription factors) that actuate them. In contrast to the prevailing MPRA approaches, we evaluated potential *cis*-regulatory elements in a relevant cell type and therapeutic vector, enabling direct clinical translation. We demonstrate that enhancer function in HUDEP- 2, but not in K562 cells, corresponded with high fidelity to their performance in adult CD34^+^-derived erythroid progenitors, highlighting the importance of using an appropriate cell context for evaluation.

We show that these new enhancer elements display precise spatiotemporal activity patterns both *in vitro* and *in vivo* as their activity is restricted to the erythroid lineage and further confined to specific stages of erythroid maturation. In the case of hemoglobinopathies current gene therapy paradigms entail the ex vivo transduction of a patient’s hematopoietic stem and progenitor cells which maintain multilineage differentiation capacity. Therefore, a constitutive promoter-enhancer pair would lead to ectopic pan-hematopoietic expression with unknown effects. Indeed, it was recently shown that the HS2 component of the μLCR enhancer has promiscuous cell type activity and thus only partial lineage specific LCR activity^30^. As such, we considered lineage specificity of these elements as an additional safety layer for these vectors.

In order to explore the potency of a μLCR–free vector design, we designed a clinically relevant vector featuring a therapeutic transgene, a truncated β-globin promoter, novel erythroid-specific enhancer elements and the C1 chromatin insulator^14^. We show that replacement of the μLCR by the *PVT1* enhancer element resulted in overall comparable potency in terms of HbF+% cells, γ-globin output, oxidative stress reduction, erythroid cell enucleation and phenotypic correction. Importantly, owing to its small size (∼10- fold shorter than the μLCR), we were able to achieve up to 5-fold increase in lentiviral particle yield in addition to significantly increased transducibility of HSPCs. These features are key to enabling the clinical-scale, GMP-grade vector production at lower MOIs and the successful transgene delivery which in turn is expected to reduce the overall commercial cost and improve patient’s accessibility^41^.

In summary, we present an integrated large-scale FACS- and lentiviral-based screening assay informed by gene regulatory atlases for the discovery of short, cell-type selective transcriptional enhancers with a broad range of amplitude. These enhancer elements can be specifically optimized for the design of safer, more efficient and lower cost production gene therapy vectors.

## Data availability

*Ex vivo* CD34^+^ erythroid differentiation DNase I data was retrieved from GEO with accession numbers GSM5554291-GSM5554326. DNase I data from CD4^+^ activated T cells, CD8^+^ activated T cells, macrophages, CD14^+^ monocytes, NK cells, CD19^+^ B cells K562 were obtained from ENCODE (accession numbers: ENCSR438USP, ENCSR294YEJ, ENCSR721XAP, ENCSR407WGG, ENCSR241BNZ, ENCSR381PXW, and ENCSR000EKS). DNase I data from HUDEP-2 cells is deposited in GEO with accession number: [DNase I from HUDEP-2 is currently in GEO submission and is available immediately by request]

## Author contributions

N.P., J.V. and G.S. designed and conceptualized the study; N.P., G.G., P.S., K.P., M.I., J.B., T.U., H.W., A.K., NI.V., MS.W. performed experiments. N.P., G.G., and J.V. analyzed data. G.S. and J.V. supervised the study. J.A.S, E.Y. and T.P. provided consultation. N.P., G.G., P.S., and J.V. wrote the manuscript.

## Acknowledgements

This work was supported by the NIH grant 5R01HL136375-02 (to G.S. and T.P.) and by an EHA Research Grant award granted by the European Hematology Association (to N.P.). This work is dedicated to the memory of Prof. George Stamatoyannopoulos who conceptualized and supervised the project until his death in June 2018.

## Methods

### Cell culture of immortalized cell lines

K562 cells were maintained at a density of 2.5x10^5^ per mL inRPMI-1640 (Thermo Fisher Scientific) with 10% heat-inactivated Fetal Bovine Serum (Thermo Fisher Scientific) and 100 units/mL streptomycin/penicillin (Life Technologies). Polybrene (8 ug/ml) was added to the culture during lentiviral transduction.

HUDEP-2 cells were cultured as previously described^42^. Briefly, during their expansion phase the cells were cultured in StemSpan H3000 medium (Stem Cell Technologies) supplemented with 100 ng/mL stem- cell factor (SCF, Peprotech), 3 IU/mL erythropoietin (EPO, Peprotech), 10−6 M dexamethasone (Sigma), 1 μg/mL doxycycline (Sigma), and 100 units/ml penicillin/streptomycin. Polybrene (8 ug/ml) was also added to the culture during lentiviral transduction. During differentiation, HUDEP-2 were maintained in Iscove’s modified Dulbecco’s medium (IMDM) supplemented with 330 μg/mL human holo-transferrin (Sigma), 10 μg/mL recombinant human insulin (Sigma), 2 IU/mL heparin (Sigma), 5% human AB plasma (obtained from Bloodworks Northwest, Seattle, WA), 3 IU/mL EPO, 100 ng/mL SCF, 1 μg/mL doxycycline (Sigma), and 100 units/ml penicillin/streptomycin.

### Cell culture of primary CD34^+^ cells

CD34^+^ cells from mobilized adult healthy donors were maintained in StemSpan H3000 with penicillin/streptomycin and the following cytokine cocktail: Flt3 ligand, thrombopoietin (TPO), and SCF (CC110, Stemcell Technologies). For lentiviral transduction the cells were plated on retronectin coated plates (Takara Bio) and transduced in the presence of 8μg/ml protamine sulfate (Sigma). Erythroid differentiation of CD34^+^ cells was carried out as previously described^43^ in 3 stages. During Stage I the cells were expanded in the presence of Hydrocortisone, IL-3, SCF and EPO, during stage II only in the presence of SCF and EPO and in the final stage III, SCF was omitted. Granulocytic/monocytic differentiation of CD34^+^ cells was achieved by culturing the cells in the same basal medium used for the *ex vivo* erythroid differentiation, with a different cytokine cocktail: SCF (500 µg/mL), IL-3 (100 µg/mL), GM-CSF (50 µg/mL), G-CSF (50 µg/mL) as previously described^44^. Megakaryocytic differentiation was performed as previously described (Georgolopoulos et al, 2021). Briefly, CD34+ HSPCs were differentiated to the megakaryocyte lineage by culturing for 11 days in IMDM based media containing 30ng/mL rhTPO (PeproTech), 1ng/mL rhSCF (PeproTech), 7.5ng/mL rhIL-6 (PeproTech), 13.5ng/mL rhIL-9 (PeproTech), 20% BIT (StemCell Technologies), 40µg/mL LDL (Millipore-Sigma), 0.05mM beta mercaptoethanol (Millipore-Sigma), and 1x Penicillin/Streptomycin (ThermoFisher Scientific).

### Selection of candidate DHS elements and DHS tiling strategy

The initial set of 11,805 DHS elements with dynamic accessibility during human *ex vivo* erythropoiesis as identified previously^12^ was filtered to exclude elements with >30 normalized DNase I counts per element at day 0 (undifferentiated CD34^+^ hematopoietic stem and progenitor cells). For each of the resulted 5,393 DHS we designed multiple 198bp-long tiles that span the entire DHS with variable overlap between any two adjacent tiles (50bp minimum) ensuring that one tile aligns with the center of the DHS. DHS smaller than 198bp would be represented by a single fragment equal to the DHS size, while larger DHS were tiled by at least 3 fragments. This resulted in a set of 15,000 tile sequences.

### Enhancer lentiviral-library construction and screening

A pool of 15,000 tiles was first chemically synthesized (Agilent Technologies) at 10 pmol each and then amplified in 9 cycles using Phusion DNA polymerase (New England Biolabs) from primers that added flanking overlaps to tiles allowing Infusion cloning in a screening vector. The screening vector was based on a lentiviral vector pRRLSIN.cPPT.PGK-GFP.WPRE (a gift from Didier Trono; Addgene plasmid # 12252), in which an enhancer blocker and barrier insulator element C1^14^ was introduced 36 bp into the proximal portion of the 3′ LTR; from this position, the insulator element is copied into the 5′ LTR during proviral cDNA synthesis, thus flanking the expression cassette on both sides upon integration. In addition, the original PGK promoter was replaced with a minimal β-globin promoter (chr11:5227033-5227201, hg38) and EcoRI and KpnI sites in positions 3,830 and 3,846 in the original pRRLSIN vector were deleted and re-created in 5’ position of the β-globin promoter to allow insertion of DHS tiles. the titer of the lentiviral library was measured using qPCR in genomic DNA isolated from K562 cells that were infected with two known volumes of lentivirus prep. qPCR was carried out using SYBR-Green protocol with primers specific to RRE region in lentiviral genome (Forward, 5’-AACTCACAGTCTGGGGCATC-3’ and Reverse, 5’-TGGTGCAAATGAGTTTTCCA-3’) and to ALB region in human genome (Forward, 5’- GCTGTCATCTCTTGTGGGCTGT-3’ and Reverse, 5’-ACTCATGGGAGCTGCTGGTTCA-3’), with a known number of DNA molecules containing RRE and ALB amplicons as a standard curve. The vast majority of designed DHS fragments (>95%) were successfully synthesized and represented in the lentiviral library (**Supplementary** Figure 3). A similar cloning strategy was used for the construction of the full- DHS library of 202 elements identified based on the 15.000 tile library. The elements were synthesized as 301-bp fragments (Twist Biosciences), pooled, and amplified using their flanking sequences with primers that added flanking overlaps to allow Infusion cloning in the screening vector. Cloning product for both libraries was introduced in chemically competent E. coli and resulting plasmid produced with redundancy of 144 clones per insert for the large tile library and 550 clones per insert for the small library. Forty candidate DHS fragments were also cloned individually in the screening vector, using identical approach, as were enhancer controls consisting of the β-globin LCR HS2 (chr11:5280523-5281381, hg38) and β- globin µLCR (chr11:5280522-5281209; 5284350-5285412; 5287870 -5288668, hg38), and a negative control consisting of sequence in ALB locus (chr4: 73,418,934-73,419,006, hg38). HUDEP-2 cells were transduced with either lentiviral enhancer screening library at MOI=0.4. 5 days post transduction the cells were sorted based on their GFP expression on a MoFlo Astrios Cell Sorter. Specifically, 3 bins were selected represented the 5% of top, medium, and low GFP expressing cells with an average MFI of 15.000, 3.000 and 750 respectively (**Supplementary** Figure 3).

### Sequencing of enhancer screening libraries and data preprocessing

Total DNA from enhancer screening library transduced HUDEP-2 cells was isolated using the DNeasy Blood & Tissue kit (Qiagen, 69504) and the enhancer inserts were PCR amplified using the following primers:

- F: 5’-ACACGACGCTCTTCCGATCTNNNNGTCGAATTAAGGACCGGATCA-3’
- R: 5’- GACGTGTGCTCTTCCGATCTNNNNAAGTGATGACAGCCGTACCA-3’

Samples were sequenced on HiSeq 4000 (Illumina) using a 2x76 paired-end sequencing. Cloning region flanking sequences were trimmed using seqtk (https://github.com/lh3/seqtk) using the *trimfq* command and removing 22 bases from read1 and 20 from read2. Illumina adapters were trimmed using Trimmomatic v0.36 setting the ILLUMINACLIP argument to 2:30:10:2:true. Sequences were aligned against the GRCh38 reference genome with BWA^45^ and sequences were filtered using samtools (http://www.htslib.org)^46^ with the following arguments: view -q 1 -f 64. Sequences were converted to genomic coordinates BED file using the *bam2bed* tool from the BEDOPS suite^47^. For every reference sequence (tile or full DHS depending on screening vector library) we counted the number of reads in each sample that overlap the reference sequence by ≥90% using the *bedmap* tool from BEDOPS.

### Enhancer screening analysis

To estimate the effect of each tile from the 15,000-tile screening library on GFP expression we employed a maximum likelihood estimation framework adapted from MAUDE^16^ developed for sorting-based screenings. The log-likelihood of the observed read counts in each GFP bin were calculated under a negative binomial model based on the expected reads per bin following a normal distribution null model. The expected mean effect score was the basis for ranking tiles for enhancer activity. For the mini-library of 203 full-size DHS sequences the effect size of each sequence on GFP expression was estimated using linear regression of the per element counts against the MFI of each GPF sorting bin.

### Lentivirus production

The VSV.G-pseudotyped Lentivirus vectors (LV) were produced in 293FT cells grown in DMEM and transiently transfected using calcium phosphate method with a plasmid mix containing packaging vector psPAX2, envelope pseudotyping vector pVSV-G, and library screening vector containing DHS tiles or elements, with or without shRNA cassette. Sixteen hours after transfection, the medium was changed and the supernatant collected twice, twenty-four and forty-eight hours after medium change. LV-containing supernatants were filtered through a 0.45-μm filter (Millipore). For the LV concentration we used Spin-XR UF concentrators (Corning). To estimate the viral titers, K562 cell were seeded in 48-well plate at 5x10^4^ cells/well and infected with serial dilutions of the vector stocks in the presence of 8 µg/mL Polybrene, maintained in culture for at least five days, and genomic DNA from cells was isolated. To estimate the physical viral titer, real-time quantitative PCR was performed in ABI 7500 (Applied Biosystems) with the use of the following primers and probes:

- Gag F: 5’-GGAGCTAGAACGATTCGCAGTTA-3’
- Gag R: 5’-GGTTGTAGCTGTCCCAGTATTTGTC-3’
- Gag Probe: 5’-FAM-ACAGCCTTCTGATGTTTCTAACAGGCCAGG-TAMRA-3’
- hAlb F: 5’-TGAAACATACGTTCCCAAAGAGTTT-3’
- hAlb R: 5’-CTCTCCTTCTCAGAAAGTGTGCATAT-3’
- hAlb Probe: 5’-VIC-TGCTGAAACATTCACCTTCCATGCAGA-TAMRA-3’

Each DNA sample was run in 25 μL reaction volume using Taqman Universal PCR Master Mix (Applied Biosystems). Thermal cycling was started for 2 min at 50°C, followed by 10 min at 95°C and 40 thermal cycles of 15 s at 95°C and 1 min at 60°C. Vector copy number/cell was calculated by normalizing to the endogenous ALB gene.

### Genome editing

To delete the two erythroid enhancers (coordinates) we designed and produced TALEN monomers, flanking in pairs the 5’ and 3’ ends of the *PVT1* and *PPARA* enhancers using adaptations of previously described methods (Cermak et al 2011, Sakuma et al. 2013). TALEN mRNAs were prepared using a mMessageMachine T7 Ultra Kit (#AM1345, Ambion). Sense and anti-sense TALENs recognize the sequences are presented in Supplementary table 3.

For all transfections, a BTX ECM830 device (BTX Harvard Apparatus) with a 2 mm gap cuvette was used. Two days post transfection we performed single cell sorting in 96 well plates. Single and double knock out clones (SKO and DKO) were evaluated by an in-out PCR of each clone with the following primers:

- PVT1 In Rev: 5’-GCCCCAGCAAAGACGTTAAG-3’
- PVT1 Out For: 5’-TGCGGAGTGAGCCTTATTCA-3’
- PVT1 Out Rev: 5’GTTGTGGGGTACAAGCCAGA-3’
- PPARA In Rev: 5’GGCCACACACATGGCTTTTT-3’
- PPARA Out For: 5’GGCCTAGGTTTTTGCTGGGA-3’
- PPARA Out Rev: 5’TTTGAGGTCATCAGCTGCGT-3’

### RNA sequencing

For gene expression analysis, total RNA from expanded HUDEP-2 edited and wild-type clones was collected using the mirVana RNA isolation kit (ThermoFisher Scientific) or RNeasy Mini Kit (Qiagen) from sorted (>20,000 cells) and bulk cultures (>1,000,000 cells). Illumina libraries were constructed using the TruSeq Stranded Total RNA with Ribo-Zero Globin (Illumina). Finally, libraries were quantified using Fragment Analyzer (Advanced Analytical). RNA-seq libraries were sequenced with HiSeq 4000 using a 2x76bp read length.

### Differential gene expression analysis

RNA-seq reads were aligned against the GRCh38 reference genome using STAR Aligner^48^. Gene counts from were obtained using featureCounts^49^. Differential expression analysis was performed using DESeq2^50^. Differentially expressed genes were called at adjusted p-value < 0.05 and absolute log2 fold change > 0.58.

### DNase I accessibility

DNase I-seq from HUDEP2 cells was performed as described previously^12^. Briefly, 100,000-200,000 live cells were collected, and nuclei were extracted in the presence of 0.04% de-ionized IGEPAL CA-630 incubated at 4 °C for 10 min. Nuclei were treated with a gradient of DNase I solution (40 IU to 100 IU of DNase I) for 3 min at 37 °C. DNase I digestion was quenched by adding an equal volume of 5X Stop buffer and 20μL RNase A (Sigma, R4642) followed by incubation at 37 °C for 60min. After incubation, 1µL of Proteinase K (Sigma, P4850) was added and the reactions were incubated at 50 °C for 60min. Digested genomic DNA was visualized on 1.2% agarose gen and the fragment size profile was generated using the Fragment Analyzer (Advanced Analytical). Prior to genomic library generation, fragments were subjected to size selection with large fragment depletion by magnetic bead separation. Fragment size distribution and concentration of the fractionated sample was measured with Fragment Analyzer (Advanced Analytical). Illumina compatible, double-stranded DNA library libraries from the size fractionated samples were constructed using the ThruPLEX DNA-seq Kit (Takara Bio) according to manufacturer’s instructions. DNase I-seq libraries were sequenced on NextSeq 500 with a 2x36bp read length. Adapter trimmed FASTQ files were aligned against GRCh38 using BWA^45^. Hotspots were detected using hotspot2 software (https://github.com/Altius/hotspot2). All downstream DNase I-seq analyses were performed on DNase I hotspots (genomic regions with statistically significant enrichment in DNase I cleavage).

### shRNA vector design and production

Lentiviral vectors were based on the pRRLSIN.cPPT.PGK-GFP.WPRE (kindly gifted from Didier Trono; Addgene plasmid # 12252; http://n2t.net/addgene:12252; RRID:Addgene 12252) in which the original human PGK promoter was exchanged for a truncated human beta-globin promoter (human genome hg38 coordinates: chr11:5,284,251-5,285,452). In addition, the vector contained a C1 insulator/barrier element^14,51^ inserted in the proximal portion of the 3′ LTR. From this position, the element is copied into the 5′ LTR during proviral cDNA synthesis, thus flanking the entire proviral expression cassette. EcoRI and KpnI sites in the original vector were deleted and recreated in the 5’ position proximal of beta globin promoter, serving as cloning sites for insertion of enhancer elements in this basal screening vector. Subsequently, positive control vector derivatives containing truncated HS2 element originating from human beta globin locus control region and a second one with expanded region composed of truncated HS2, HS3, and HS4 elements^52^ were created. Human genome hg38 coordinates for HS2 are chr11: 5,280,523-5,281,381, hg38, coordinates for HS2-HS3-HS4 are chr11: 5,280,522-5,281,209, 5,284,350-5,285,412, 5,287,870-5,288,668, respectively. Negative control vector was prepared by inserting a sequence from human albumin locus (human hg38 coordinates chr4: 73,418,934-73,419,006). Lentiviral vectors expressing miRNA-embedded shRNA targeting *BCL11A* were based on control vectors described above in which the GFP ORF was appended by a cassette containing *BCL11A*-targeting shRNA embedded in miR223 as described previously^17,18^.

### FACS

To assess the multilineage engraftment of CD34^+^ cells in the bone marrow of NBSGW mice post transplantation, the following antibodies were used: CD45-APC (BD Biosciences), CD19-PE (BD Biosciences), CD3-PE (ExBIO), CD33-PerCp (ExBIO), CD235a-PE (ExBIO), CD34(), CD41. After wash, cells were resuspended in FACS buffer and analyzed using a FACS-Calibur (BD Biosciences, San Jose, CA). To measure the percentage of HSPCs in peripheral blood of NBSGW mice post mobilization, the following antibodies were used: murine HSPCs: mouse Lineage Cocktail-APC, CD117-PE, LY-6A/E (sca- 1)-PerCP (BD Biosciences, San Jose, CA); human HSPCs: CD45-PerCP, CD34-APC, CD46-FITC (BD Biosciences, San Jose, CA). Debris was excluded using a forward scatter-area and sideward scatter-area gate. Flow cytometry data were then analyzed using FlowJo (version 10.0.8, FlowJo, LLC).

### Globin HPLC

Individual globin chain levels from transduced *ex vivo* differentiated normal CD34^+^ cells or NBSGW bone marrow-engrafted human cells were quantified on a Shimadzu Prominence instrument with an SPD-10AV diode array detector and an LC-10AT binary pump (Shimadzu, Kyoto, Japan). Vydac 214TP™ C4 Reversed-Phase columns for polypeptides (214TP54 Column, C4, 300 Å, 5 µm, 4.6 mm i.d. x 250 mm) (Hichrom, UK) were used. Globin chains from *ex vivo* differentiated thalassemic cells were quantified on a Shimadzu LC-2060C 3D Liquid chromatography with a GmbH MultoHigh Bio 300, 250x3 mm column. A 38%-60% gradient mixture of 0.1% trifluoroacetic acid in water/acetonitrile was applied at a rate of 1 mL/min.

### Xenotransplantation

Immunodeficient NOD.Cg-KitW-41J Tyr^+^ Prkdcscid Il2rgtm1Wjl/ThomJ (NBSGW) mice were obtained from the Jackson Laboratory (Bar Harbor, ME). To evaluate the enhancers’ cell specificity, CD34+ cells were transduced with a pool of the same MOI of 40 different vectors (**Figure 2**). 2 days post transduction, *ex vivo* transduced 1x10^6^ CD34^+^ cells from healthy donors were injected intravenously into NBSGW recipient mice, without any conditioning. Non-transduced CD34^+^ cells were used as controls. Sixteen weeks post transplantation, NBSGW mice were sacrificed and bone marrow cells were collected, for assessment of multilineage engraftment. To evaluate specificity, efficiency and *in vivo* safety of the *BCL11A*-shRNA vectors, CD34^+^ cells from healthy donors were transduced with the same MOI (40) with the two vectors. Transplantation of the cells was performed in the same manner as described above. Sixteen weeks post transplantation the recipient mice were sacrificed, and bone marrow was collected from 4 long bones. Multilineage engraftment, GFP and HbF expression was evaluated by flow cytometry.

### Cytospin slide preparation

Cytospins of 0.3-0.5x10^5^ thalassemic transduced and untransduced cells were prepared during erythroid differentiation, by cytocentrifugation (ROTOFIX 32, Hettich Zentrifugen) at 500 rpm for 5 minutes. Cytospins were air dried and then stained with Giemsa/May-Grünwald (Merck, Darmstadt, Germany) for 8 and 3 minutes, respectively.

### Analysis of Reactive Oxygen Species (ROS) levels

Intracellular ROS levels from erythroid cells were determined using the CellROX Deep Red Flow Cytometry Assay kit (Invitrogen by Thermo Scientific), according to the manufacturer’s instructions. Briefly, 1x10^6^ thalassemic *ex vivo* differentiated cells were incubated for 30 minutes at 37°C, protected from exposure to light with the CellROX reagent at a final concentration of 500nM.Oxidation of the probe can be detected by the increase of fluorescence (PE) with flow cytometry.

## Supplementary Figure Legends

***Supplementary Figure 1.Features of selected DHSs active during human ex vivo erythropoiesis.***

**(a)** Erythroid differentiation day where shortlisted DHS exhibit maximum accessibility. **(b)** Upset plot with genic features overlapped by the selected DHS. **(c)** Normalized density of DNase I accessibility around ±5kb around the 5,393 selected DHSs across several non-erythroid primary cell types and immortalized cell lines.

***Supplementary Figure 2.Converting DHSs into a library of fixed size tiles.***

**(a)** Histogram of DHS size distribution with median size highlighted. **(b)** Frequency barplot depicting the number of tiles DHS are represented by.

***Supplementary Figure 3.Correspondance between DHS accessibility profiles between ex vivo generated erythroid progenitors, HUDEP-2 and K562 cell lines.***

**(a)** Relationship between DNase I density of the selected 5,393 DHS observed in day 7 *ex vivo* erythroid progenitors against K562, and HUDEP-2. Adjusted R-squared is shown.

***Supplementary Figure 4. A FACS sorting scheme to identify tile effects on GFP expression.***

**(a)** Gating strategy applied for sorting based on GFP intensity **(b)** Three equiproportional bins (low, medium, and high for GFP) each consisting of 5% of total live cells (top) and validation GFP intensity histograms of the 3 selected bins measured after FACS sorting.

***Supplementary Figure 5. Relationship between library coverage and replicate concordance.***

**(a)** Scatterplots depicting correlation (Pearson’s *r*) between replicates at various library coverage levels. **(b)** Percent (mean from *n*=3 experiments) of tiles recovered across GFP gated populations at different coverage levels. **(c)** Relationship between median reads per tile (counts) and replicate correlation (Pearson’s *r*) at 100X, 200X, and 800X coverage, respectively. Mean±SE from *n*=3 experiments is shown.

***Supplementary Figure 6. GFP expression effect correspondence between tiles and their respective full-size DHSs.***

Correlation (Pearson’s r) between the effects on GFP expression of full-size DHS sequences (*x*-axis) and the strongest tile of the corresponding original DHS sequence (*y*-axis). Pearson’s *r* correlation is shown.

***Supplementary Figure 7. Comparison between sequencing based estimate of enhancer activity and observed GFP intensity.***

Scatterplot depicting the correlation (Pearson’s *r*) between the estimated effect of individual inserts on GFP from the mini-library pooled approach, and their matched GFP MFI readings post transduction in HUDEP- 2 cells (average of *n*=3 experiments).

***Supplementary Figure 8. Identifying the gene targets of two identified enhancer elements.***

**(a)** Map of the binding positions on hg38 reference genome of TALEN dimers used to delete *PVT1* and *PPARA* enhancer elements. (**b)** Manhattan plots showing the top differentially expressed genes between *PVT1* enhancer (left) and *PPARA* enhancer (right) vs. mock-transduced control. *x*-axis is the gene position on genome and *y*-axis is the –log_10_ adjusted *p*-value. Points are colored by log_2_ fold change.

***Supplementary Figure 9. Correlation of vector MOI and percentage of transduced cells.***

**(a)** Scatterplot demonstrating the relationship between vector MOI and resulting fraction of GFP^+^ cells for each of the three vectors. Lines are regression fitted trends. **(b)** Barplots showing the differences between the percentage of transduced cells and different MOI. (c) Differences in percent HbF+ cells within the GFP+ population between different vectors, across three CD34+ donors. Likelihood ratio test p-values are shown testing for changes across MOI values (n=3). (d). Differences in percent of HbF+ positive cells within the GFP+ population between different MOI values across three CD34+ donors. T-test p-values shown (n=3)

