## Supplemental Figure 1 for "Large-scale discovery of potent, compact and lineage specific enhancers for gene therapy vectors"

### Supplementary Figure 1

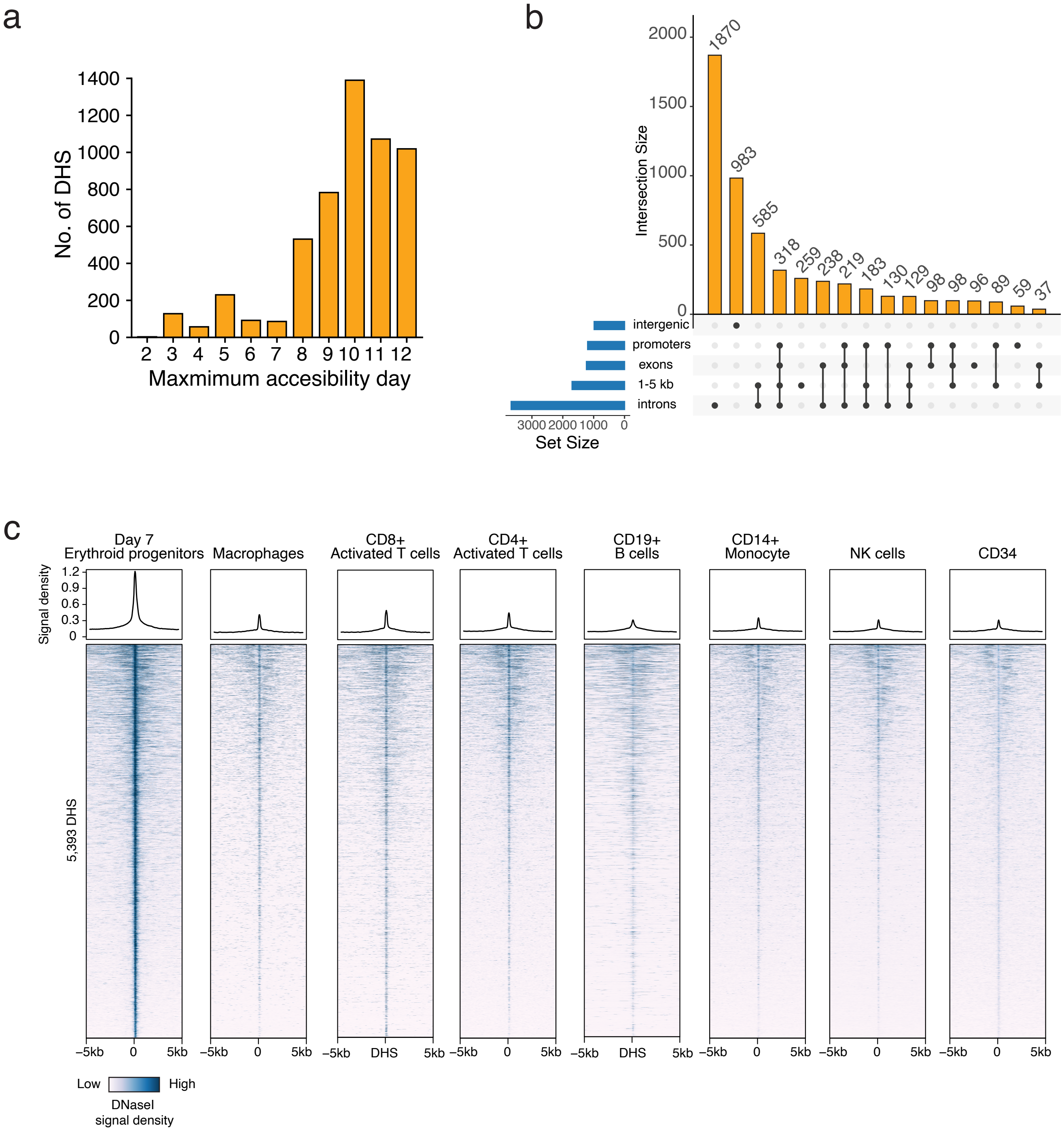


**Supplementary Figure 1.** ***Features of selected DHSs active during human ex vivo erythropoiesis.*** (a) Erythroid differentiation day where shortlisted DHS exhibit maximum accessibility. (b) Upset plot with genic features overlapped by the selected DHS. (c) Normalized density of DNase I accessibility around ±5kb around the 5,393 selected DHSs across several non-erythroid primary cell types and immortalized cell lines.
