## Supplemental Figure 2 for "Large-scale discovery of potent, compact and lineage specific enhancers for gene therapy vectors"

### Supplementary Figure 2

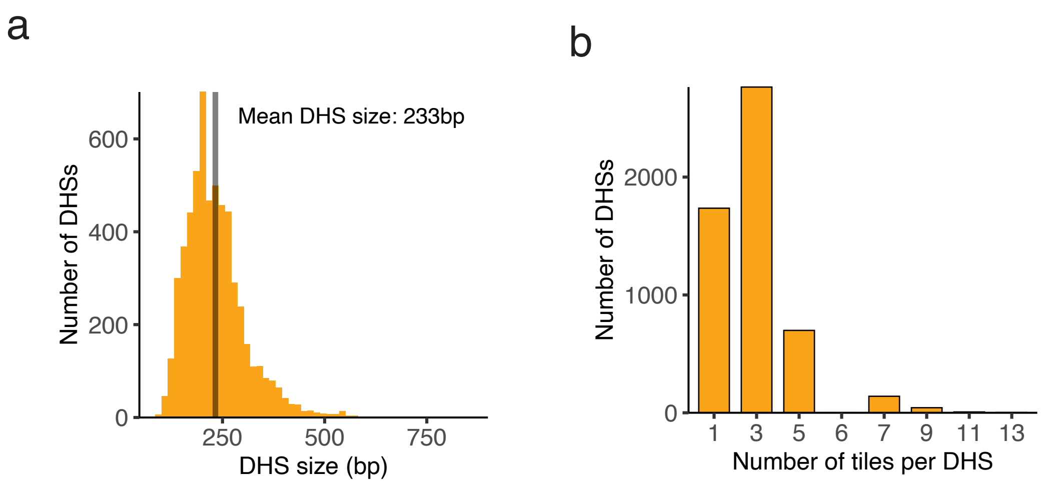


**Supplementary Figure 2.** ***Converting DHSs into a library of fixed size tiles.***
(a) Histogram of DHS size distribution with median size highlighted. (b) Frequency barplot depicting the number of tiles DHS are represented by.
