## Supplemental Figure 3 for "Large-scale discovery of potent, compact and lineage specific enhancers for gene therapy vectors"

### Supplementary Figure 3

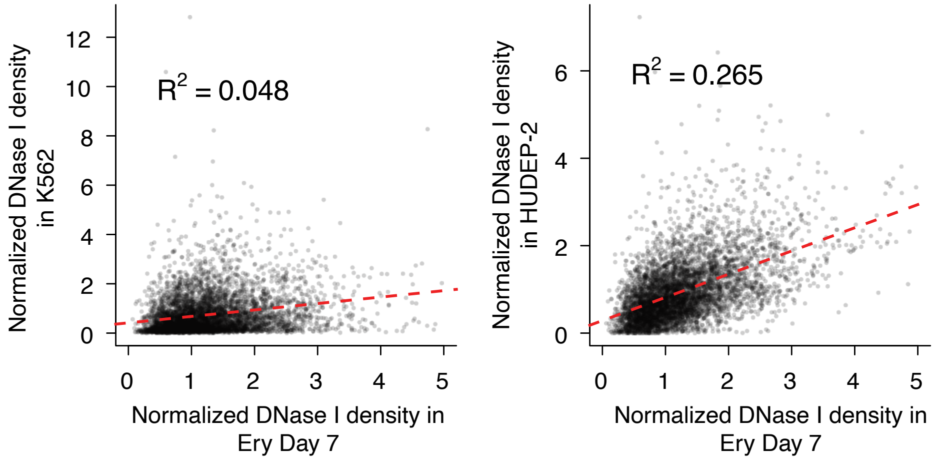


**Supplementary Figure 3.** ***Correspondance between DHS accessibility profiles between ex vivo generated erythroid progenitors, HUDEP-2 and K562 cell lines.*** (a) Relationship between DNase I density of the selected 5,393 DHS observed in day 7 ex vivo erythroid progenitors against K562, and HUDEP-2. Adjusted R-squared is shown.
