## Supplemental Figure 4 for "Large-scale discovery of potent, compact and lineage specific enhancers for gene therapy vectors"

### Supplementary Figure 4

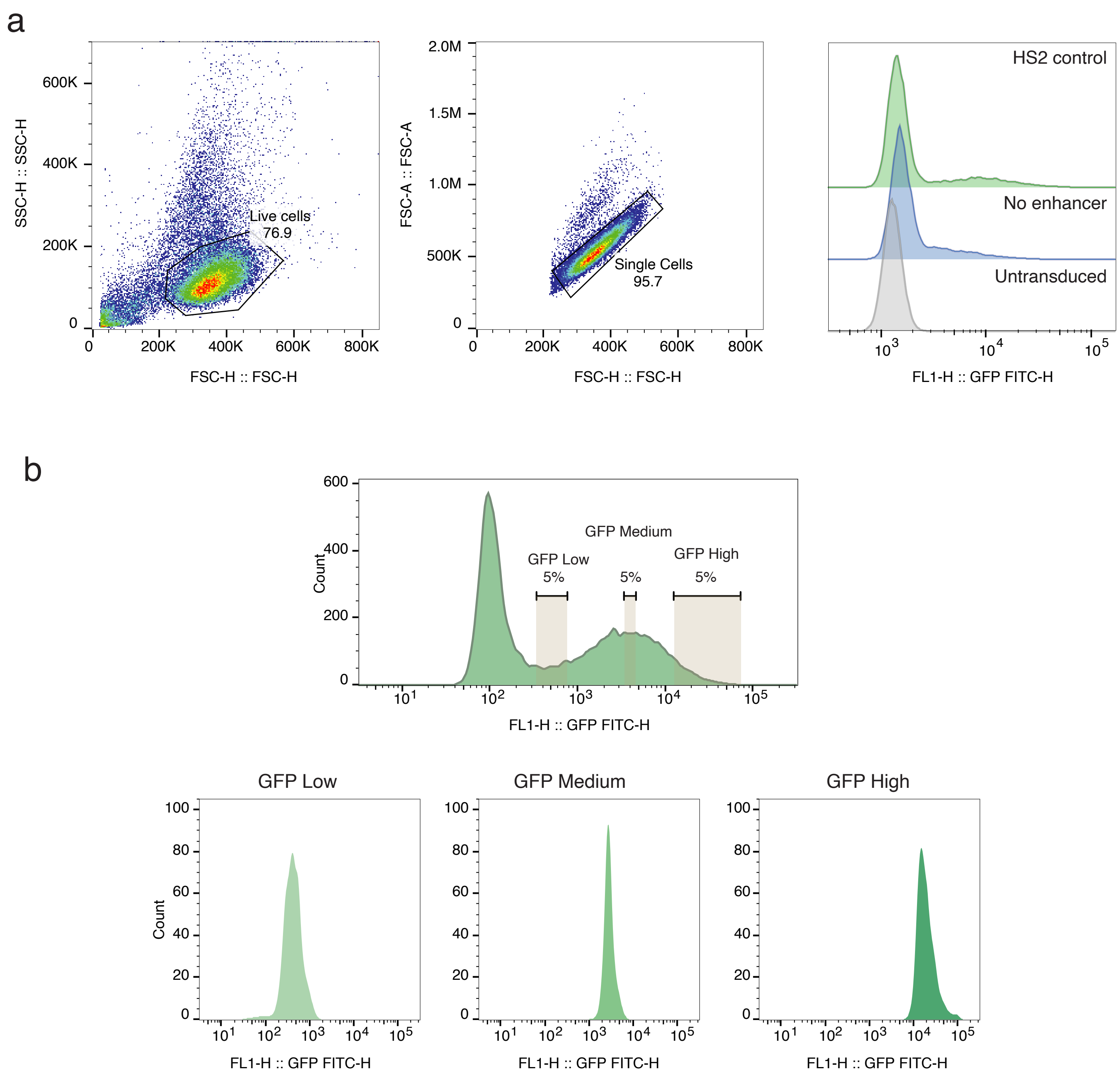


**Supplementary Figure 4.** ***A FACS sorting scheme to identify tile effects on GFP expression.***
(a) Gating strategy applied for sorting based on GFP intensity (b) Three equiproportional bins (low, medium, and high for GFP) each consisting of 5% of total live cells (top) and validation GFP intensity histograms of the 3 selected bins measured after FACS sorting.
