## Supplemental Figure 5 for "Large-scale discovery of potent, compact and lineage specific enhancers for gene therapy vectors"

### Supplementary Figure 5

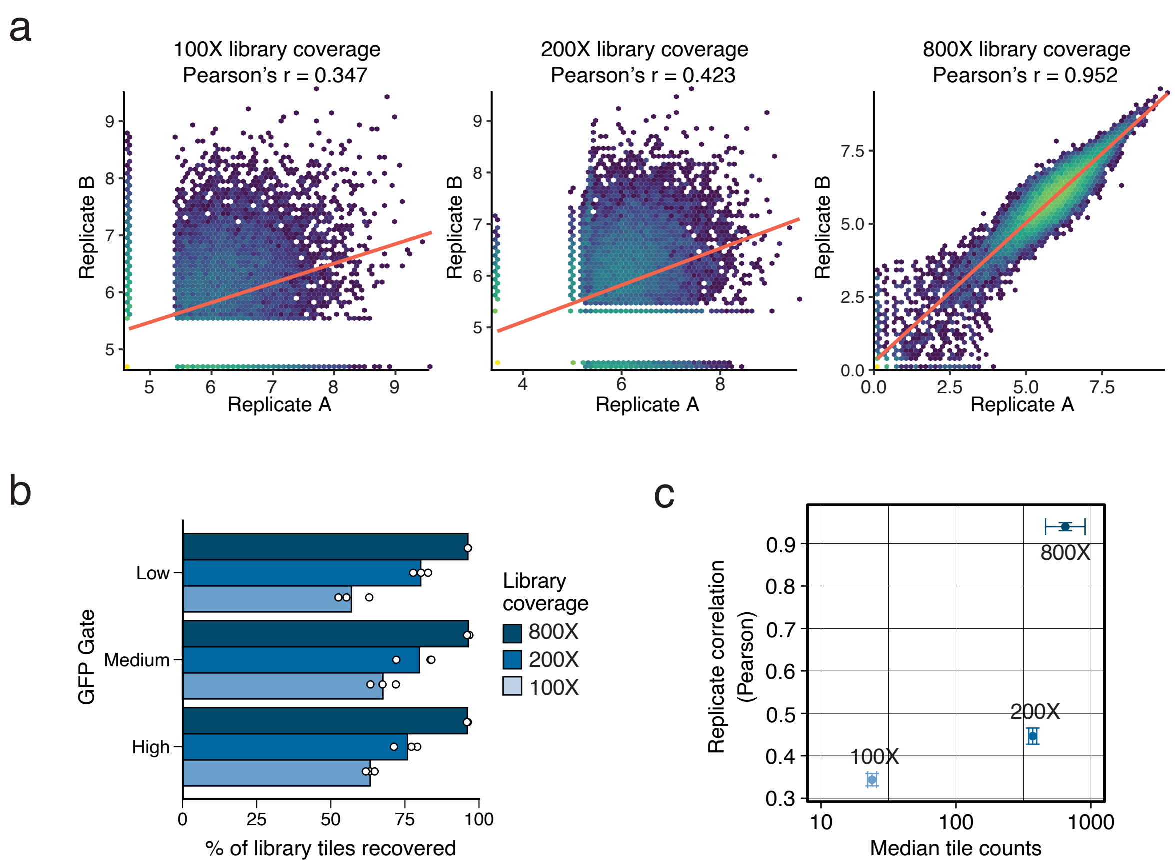


**Supplementary Figure 5. *Relationship between library coverage and replicate concordance*.**
(a) Scatterplots depicting correlation (Pearson's r) between replicates at various library coverage levels. (b) Percent (mean from n=3 experiments) of tiles recovered across GFP gated populations at different coverage levels. (c) Relationship between median reads per tile (counts) and replicate correlation (Pearson's r) at 100X, 200X, and 800X coverage, respectively. Mean±SE from n=3 experiments is shown.
