## Supplemental Figure 6 for "Large-scale discovery of potent, compact and lineage specific enhancers for gene therapy vectors"

### Supplementary Figure 6

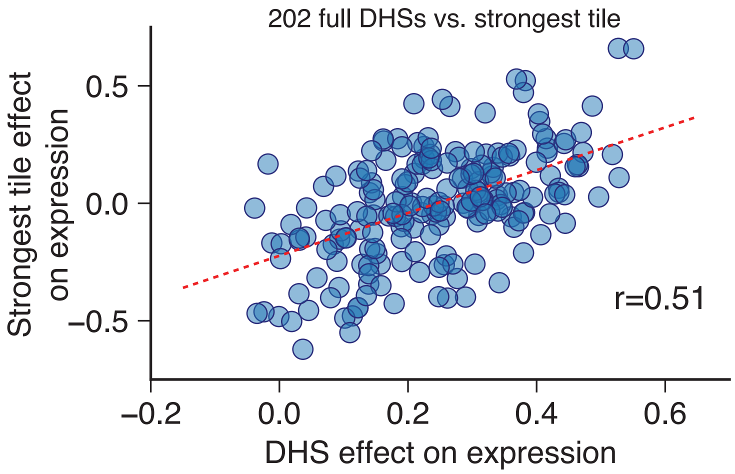


**Supplementary Figure 6. *GFP expression effect correspondence between tiles and their respective full-size DHSs.***

Correlation (Pearson's r) between the effects on GFP expression of full-size DHS sequences (x-axis) and the strongest tile of the corresponding original DHS sequence (y-axis). Pearson's r correlation is shown.
