## Supplemental Figure 7 for "Large-scale discovery of potent, compact and lineage specific enhancers for gene therapy vectors"

### Supplementary Figure 7

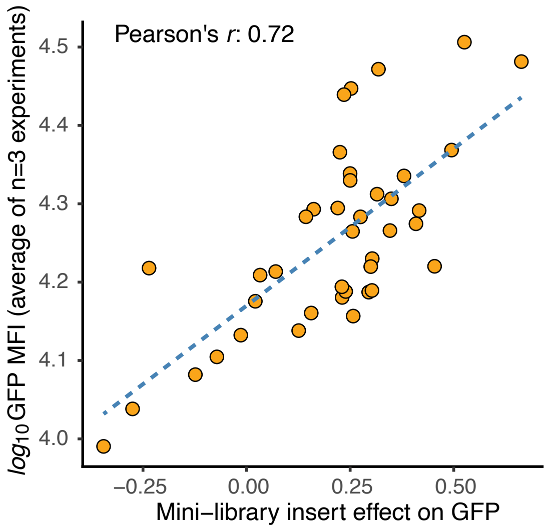


**Supplementary Figure 7.** ***Comparison between sequencing based estimate of enhancer activity and observed GFP intensity.*** Scatterplot depicting the correlation (Pearson's r) between the estimated effect of individual inserts on GFP from the mini-library pooled approach, and their matched GFP MFI readings post transduction in HUDEP-2 cells (average of n=3 experiments).
