## Supplemental Figure 8 for "Large-scale discovery of potent, compact and lineage specific enhancers for gene therapy vectors"

### Supplementary Figure 8

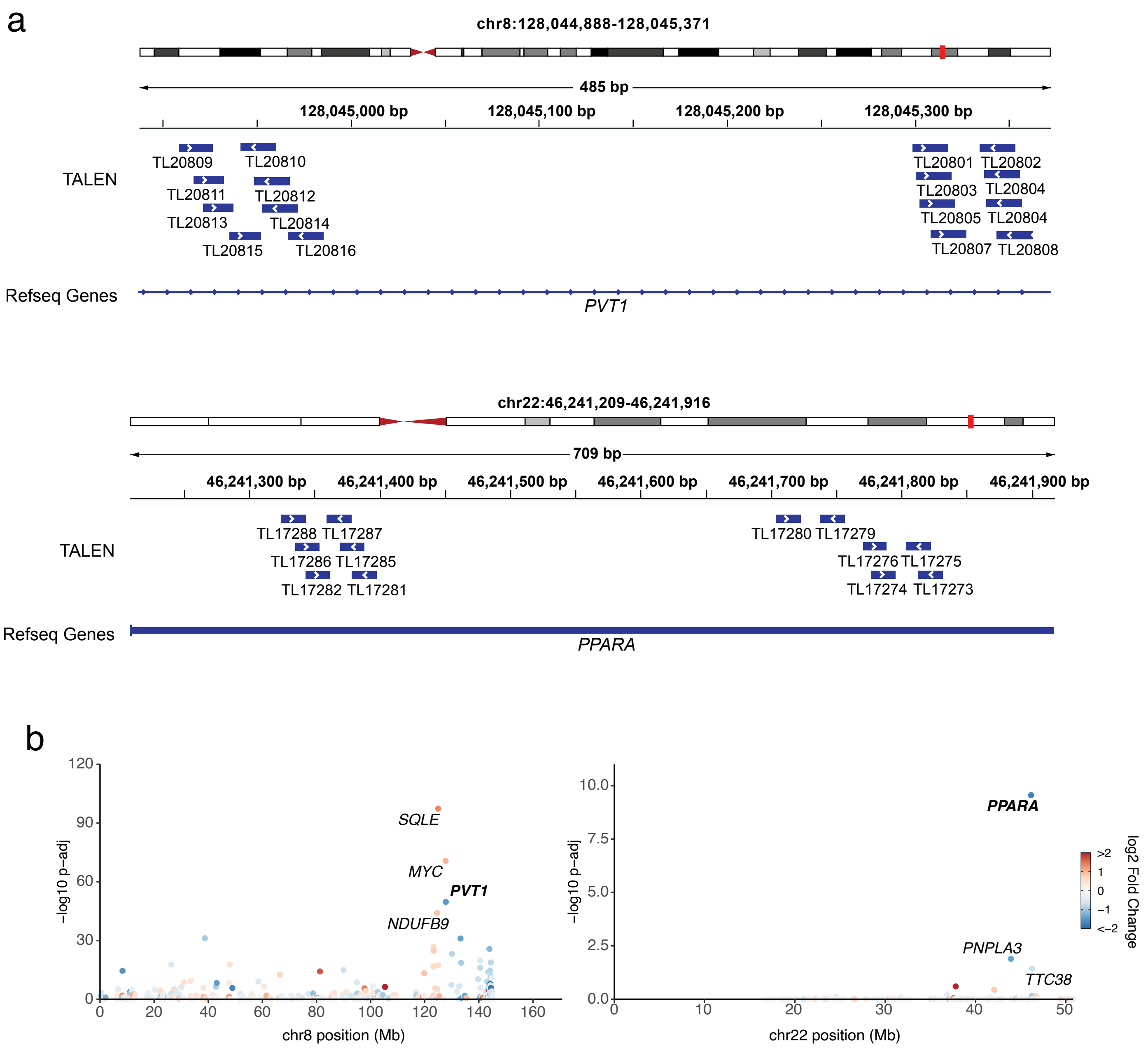


**Supplementary Figure 8.** ***Identifying the gene targets of two identified enhancer elements.***
(a) Map of the binding positions on hg38 reference genome of TALEN dimers used to delete PVT1 and PPARA enhancer elements. (b) Manhattan plots showing the top differentially expressed genes between PVT1 enhancer (left) and PPARA enhancer (right) vs. mock-transduced control. x-axis is the gene position on genome and y-axis is the –log10 adjusted p-value. Points are colored by log2 fold change.
