## Supplemental Figure 9 for "Large-scale discovery of potent, compact and lineage specific enhancers for gene therapy vectors"

### Supplementary Figure 9

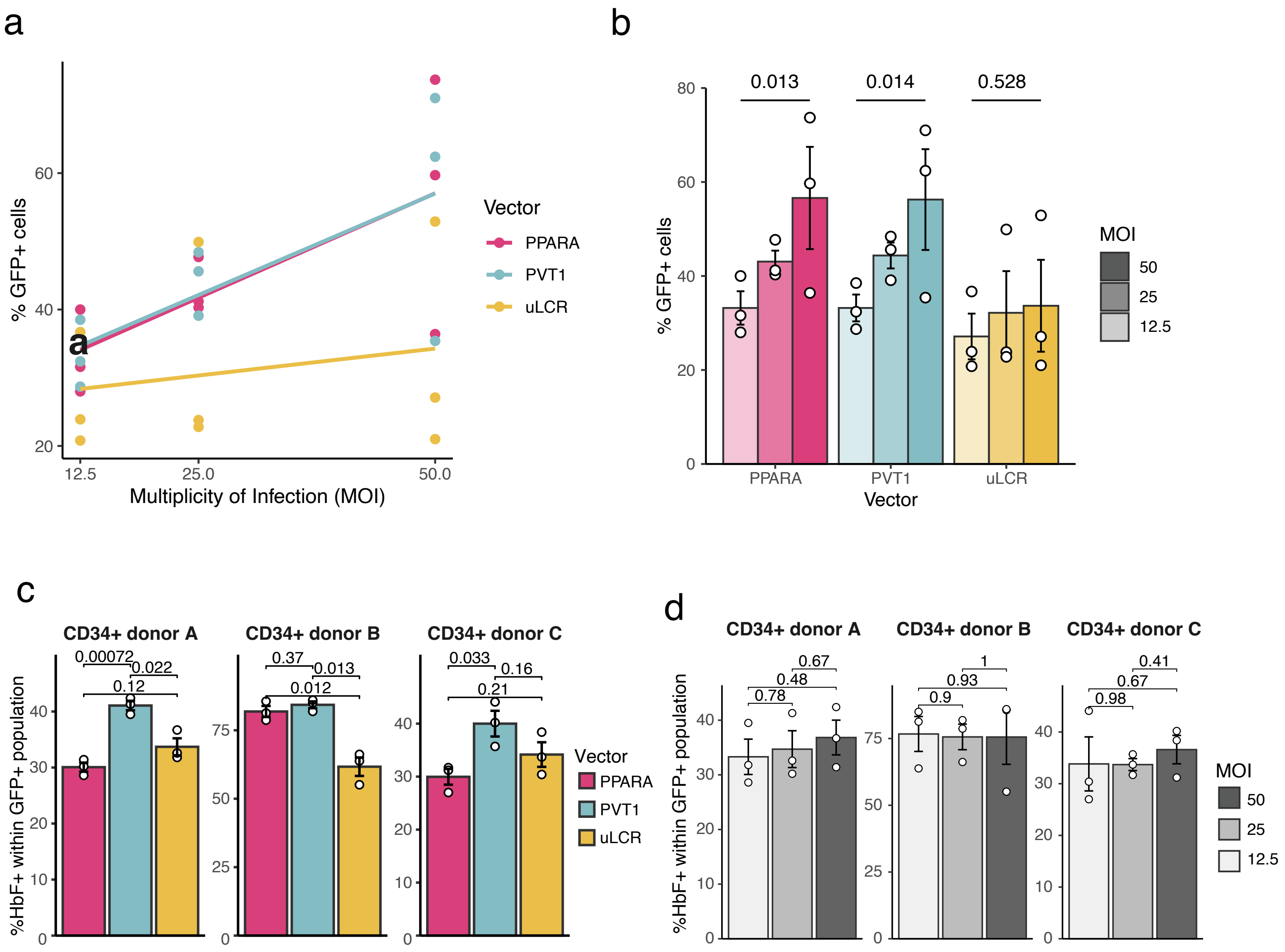


**Supplementary Figure 8.** ***Correlation of vector MOI and percentage of transduced cells.*** (a). Scatterplot demonstrating the relationship between vector MOI and resulting fraction of GFP+ cells for each of the three vectors. Lines are regression fitted trends. (b). Barplots showing the differences between percent of GFP+ cells across the three constructs. Likelihood-ratio test p-values are shown to testing whether GFP+ is a function of MOI in each vector construct. (c). Differences in percent of HbF+ cells within the GFP+ population between different vectors across three CD34+ donors. Likelihood ratio test p-values are shown testing for changes across MOI values (n=3). . (d). Differences in percent of HbF+ positive cells within the GFP+ population between different MOI values across three CD34+ donors. T-test p-values shown (n=3).
